# Distant Dipoles: A Multi-Parameter and Multi-Objective Analysis of RF Coil Performance For 7T Body MRI

**DOI:** 10.64898/2026.04.29.721770

**Authors:** Tobey D. Haluptzok, Alireza Sadeghi-Tarakameh, Russell L. Lagore, Gregory J. Metzger

**Author notes:** **CORRESPONDING AUTHOR:** Gregory J. Metzger, Center for Magnetic Resonance Research (CMRR), University of Minnesota, 2021 6th Street SE, Minneapolis, MN 55455. **FUNDING INFORMATION:** NIH P41 EB027061, NIH R01 EB029985.

## Abstract

**Purpose:** To address the limitations of single-distance, 1D performance metrics in RF coil design. This work introduces a multi-objective, volume-of-interest (VOI) based analysis to systematically characterize the trade-offs between power efficiency, pSAR efficiency, and homogeneity as a function of dipole length (*l*) and distance-to-load (*d*) for multiple dipole geometries and target anatomies.

**Methods:** Electromagnetic simulations of straight and end-meandered dipole antennas were performed with varying lengths (100-500 mm) and distance-to-load (1-81 mm) over three anatomical targets (prostate, kidney, heart). Homogeneity, power efficiency, pSAR efficiency, and load sensitivity performance metrics were calculated within each anatomical VOI. Inter-element coupling at variable *d* was assessed in a 3-element array, and a subset of single-element simulations was experimentally validated using B_1_^+^ mapping.

**Results:** A fundamental trade-off was found between power efficiency and pSAR efficiency. Optimal power efficiency was achieved with shorter dipoles (150 mm < *l* < 300 mm) closer to the sample (*d* < 30 mm), while optimal pSAR efficiency and homogeneity were achieved with longer dipoles at further from the sample (*d* > 60 mm). Inter-element coupling increased with distance-to-load but could be managed by increasing element spacing. Experimental measurements were in good agreement with simulation trends.

**Conclusion:** Increasing distance-to-load to 40-60 mm, compared with commonly used distances of 20-30 mm, offers a practical strategy for improving pSAR efficiency and homogeneity with a minimal decrease in power efficiency. This work provides a quantitative analysis that enables RF coil designers to make informed, data-driven decisions when developing next-generation body arrays and suggests that unshielded end-meandered dipoles could be an optimal transmit element geometry.

## INTRODUCTION

In 2017, the first 7T MRI system received FDA approval for clinical head and knee imaging, with developments at this higher field being driven by the promise of improved signal-to-noise ratio (SNR)^1–4^, and more favorable contrast mechanisms for some applications^5–10^. However, clinical body imaging at 7T remains investigational due to persistent difficulties in imaging reliability stemming from challenges associated with the short in-vivo wavelength. The first challenge is inhomogeneous B_1_^+^ which results in variable flip angles and, consequently, inconsistent signal and contrast across the imaging volume. The second challenge is a high peak local specific absorption rate (pSAR)^11,12^ which is a potential patient safety concern. Unfortunately, both of these issues are further exacerbated by the use of local transmit arrays^16,17^, which are ubiquitous across 7T systems^18–22^.

To mitigate these issues, dipole antennas have become an essential element for 7T body arrays, as their curl-free current patterns result in more optimal field distributions at UHF^24–27^. Consequently, the literature is rich with dipole optimizations, including geometric modifications like meandering^28–32^, unique feeding techniques^21,33^, the integration of high-dielectric materials^34,35^, and flexible dipoles^36,37^. However, to manage the large parameter space of these investigations, performance is often characterized at a fixed distance-to-load. While some studies have evaluated the dependence of element distance on individual performance metrics^38–43^, a comprehensive analysis exploring the simultaneous trade-offs between *multiple* performance metrics as a function of distance has, to our knowledge, not been reported. Furthermore, coil performance is often quantified by reducing the volumetric B_1_^+^ field to a single-line profile for analysis^36,37,44,45^. While one-dimensional plots provide valuable insights and are straightforward to implement, they do not fully capture the performance in the complex, three-dimensional anatomical imaging use-case. The ultimate clinical utility of a coil depends not on its performance along a single line, but on its ability to generate a sufficiently strong and homogeneous field across an entire target volume.

In this study, we address these gaps by presenting a comprehensive, multi-parameter, multi-objective analysis of dipole antenna performance. We systematically investigate a large parameter space of different dipole lengths, distance-to-loads, geometries, and shielding and report on multiple metrics simultaneously to transparently characterize fundamental trade-offs in performance. Crucially, these performance metrics are calculated using a volume-of-interest (VOI) based evaluation. Instead of using 1D profiles, we calculate mean power efficiency 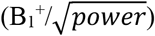, mean pSAR efficiency 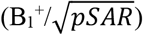, and homogeneity (mean(B_1_^+^)/SD(B_1_^+^)) directly within three-dimensional, anatomically-defined target volumes (prostate, kidney, and heart). Additionally, we evaluate how inter-element coupling and loading sensitivity are affected by these parameters, providing a more complete dataset to guide future RF coil array designs.

## Methods

An illustration of the simulation setup with all the parameters that were varied in this study is shown in Figure 1. The parameters are as follows: distance-to-load (*d*), z-shift (z), inter-element spacing (*s*), dipole length (*l*), dipole meander width (*w*), and shielded vs unshielded. All Stl coil files can be found in supplementary data.

**Figure 1.**
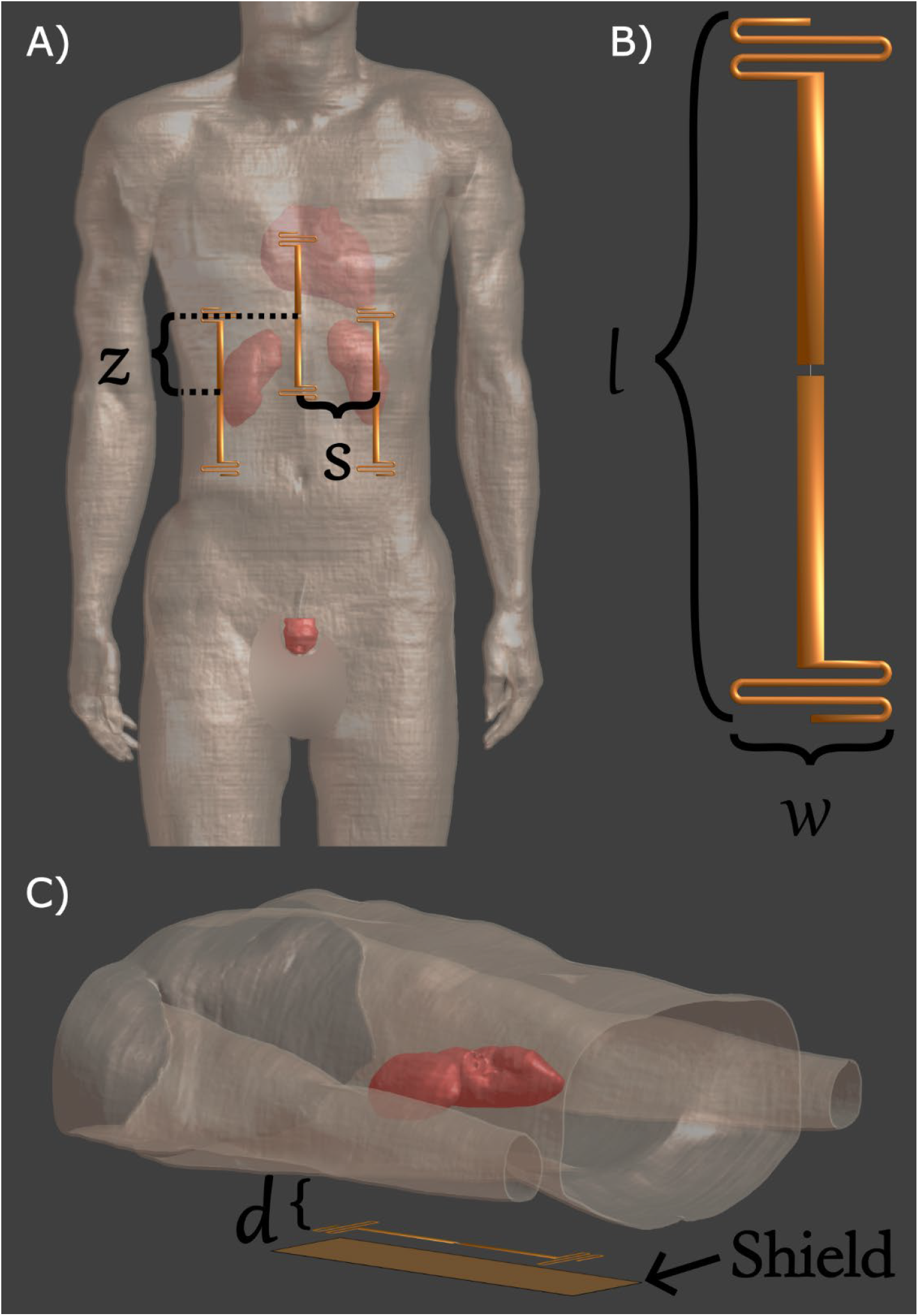
Simulation parameters varied to determine their impacts on power, pSAR, and homogeneity, sensitivity to loading, and inter-element coupling performance metrics. When the dipole meander width is set to 0, the element is a straight dipole without any meanders at the end. Abbreviations: *z*, z-shift; *s*, element spacing; *l*, dipole length; *w*, meander width; *d*, distance-to-load.

### Dipole elements and simulation

Straight dipoles, with a 4mm gap between the legs and lengths varying from 100mm to 500mm in steps of 20mm, were imported into Sim4life (Zurich Medtech, Zurich, Switzerland) for electromagnetic (EM) simulation. The feed of the dipoles was positioned over the center of the prostate, kidney, and heart of the Duke human body model^46,47^ with distance-to-load ranging from 1 to 81mm in steps of 10mm. The human body model was placed inside a perfect electric conductor cylinder of diameter 670mm to simulate the gradient shield of the MRI scanner. The elements were positioned posteriorly for the prostate and kidney simulations and anteriorly for the heart simulation. Elements were centered over the left kidney and only the fields in this kidney were analyzed. Dipoles were voxeled at 1mm resolution, the Duke model at 4mm resolution, and the gradient shield used Sim4life’s automatic voxelization settings. All simulations used a harmonic 297 MHz excitation and - 30dB convergence criterion. Electromagnetic (EM) fields were normalized to 1W accepted power, and exported to MATLAB (MathWorks, Natick, MA) for interpolation onto a 4mm isotropic grid. The E-fields, mass density, and conductivity were used to calculate 10g averaged SAR using publicly available methods^48^. The pSAR for each simulation scenario was identified as the voxel with the highest 10g SAR.

### Single Element Data Analysis

After the EM fields were post-processed, the magnitude B_1_ ^+^ field was normalized by 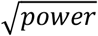 and 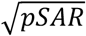 to obtain power and pSAR efficiency maps which were masked to the organ of interest in each simulation setup (i.e. the prostate, kidney, or heart). We then defined power efficiency and pSAR efficiency to be the mean normalized B_1_ ^+^ in each VOI, and also report homogeneity in these VOIs since it is an important factor for final image quality. These metrics are visualized in colormaps, with each pixel in the colormaps corresponding to the performance of a specific length and distance-to-load. The tradeoff between power efficiency and pSAR efficiency can be viewed as a multi-objective optimization problem, which is characterized in L-curves. To create these L-curves, performance for each length dipole was plotted in the pSAR efficiency-by-power efficiency space. To determine how sensitive each dipole is to variable loading, each unique length dipole was matched at a distance-to-load of 40mm via an L-match. This reference L-match was applied to the same length elements at varying distance-to-load settings and the match at each distance was recorded.

### End-Meandered Dipole Elements

To evaluate performance trends in more realistic geometries, the analysis was extended to include end-meandered dipole elements^28,29,32^ with meander widths (*w)* of 40 and 80mm. Based on the straight dipole results, the search space was narrowed to focus on the high-performance region near the knee of the L-curves consisting of lengths 220 to 300mm and distance-to-load of 20, 40, and 60mm. Simulations were conducted with and without a shield 20mm from the elements. Meandered dipoles were simulated over the heart since this anatomy had the largest performance variability in the straight dipole simulations. The same MATLAB post-processing and performance metric calculations used for the straight dipoles were also performed for the meandered dipoles. Due to the difficulty in displaying both shielded vs unshielded results in heatmaps, meandered dipole results are reported in the pSAR efficiency-by-power efficiency and CV-by-power efficiency spaces.

### Experimental measurements

To validate a subset of the numerical findings, dipoles of lengths 200, 300, and 500mm were constructed. They were tuned and matched, using a series inductor and lattice balun, to better than -18dB at a distance of 10mm and 40mm from a phantom mimicking the human body (ε_r_=51.4, σ=0.57S/m). B_1_ ^+^ maps were acquired for each element using the AFI acquisition method^49^. Line profiles were drawn perpendicular to the elements to report how the different configurations performed as a function of depth into the phantom.

### Array Simulations

The 220mm straight dipole, due to its reasonable performance and ability to accommodate a z-shift without making the overall array excessively long, was selected to be simulated in a 3-element array to investigate coupling performance. Since coupling is a complex phenomenon with many different contributing factors, this search space was not exhaustively evaluated. Instead, *s, d*, and z-shift parameters were varied. These simulations were post-processed using a custom-built MATLAB co-simulation environment that computed L-matches for each element which reduced reflection coefficients to less than -30dB and worst-case coupling was recorded for each simulation scenario.

## Results

### Single Elements

Figure 2 shows power efficiency, pSAR efficiency, and homogeneity heatmaps for straight dipoles as *distance-to-load* and *length* are varied. In each anatomy, there is an optimal power efficiency which varies between the three anatomies tested. In all cases, the optimal power efficiency occurs between *l*=150mm and *l*=300mm. In both the pSAR efficiency and homogeneity metrics, the longest dipoles at the farthest distance-to-load had the best performance but this came at a cost of decreased power efficiency. Figure 3 plots each simulation scenario’s performance in the power efficiency-by-pSAR efficiency space. The best performance in this plane is the point closest to the top right corner of the plot, which corresponds to both the highest power and pSAR efficiency. The dominating edge of these curves, indicated by colored arrows in Figure 3A, is comprised of configurations with different lengths and different distances. One definition of the “ideal” tradeoff is at the knee of the L-curve which occurs between a distance-to-load of 30mm and 60mm and a length of 200mm to 300mm.

**Figure 2.**
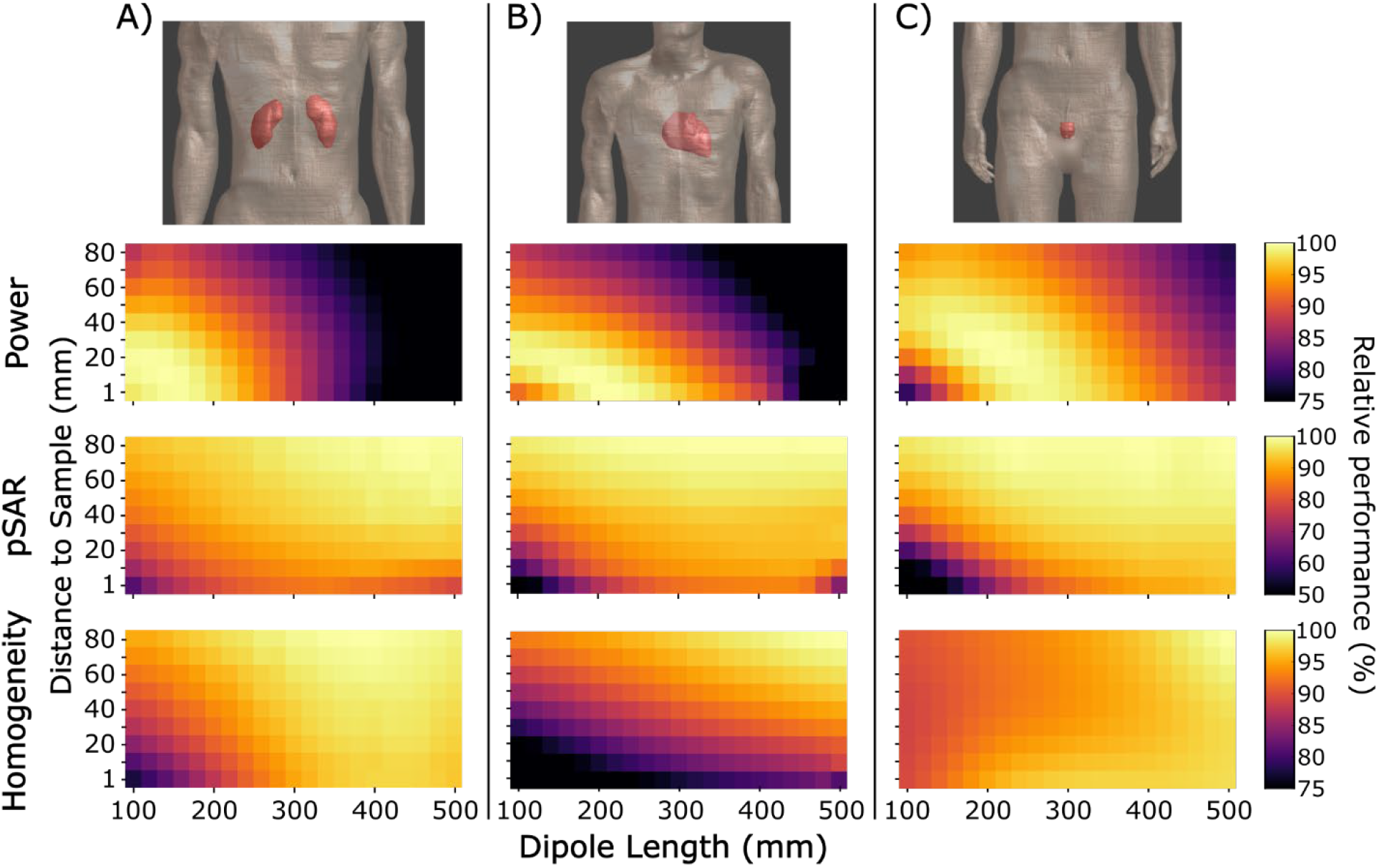
Straight dipole performance metrics when varying dipole length and distance-to-load. As the anatomy of interest moves further into the body (in increasing order of kidneys, heart, prostate). The optimal power efficiency scenario favors shorter dipoles at shorter distance-to-load. For optimal pSAR and homogeneity, the longest dipole at the furthest distance is preferred. When using these figures to determine what scenario you want to use for an array, it will be beneficial to consider the prostate power efficiency and the heart homogeneity metrics since most regions in the body will be further away for most elements in the array since these single elements were positioned directly over the anatomies of interest. Special interest should be taken in the heart’s homogeneity performance since the heart has both superficial and deeply located regions. Therefore, the heart’s homogeneity metric is a good measure for overflipping near the surface and decay rate into the sample

**Figure 3.**
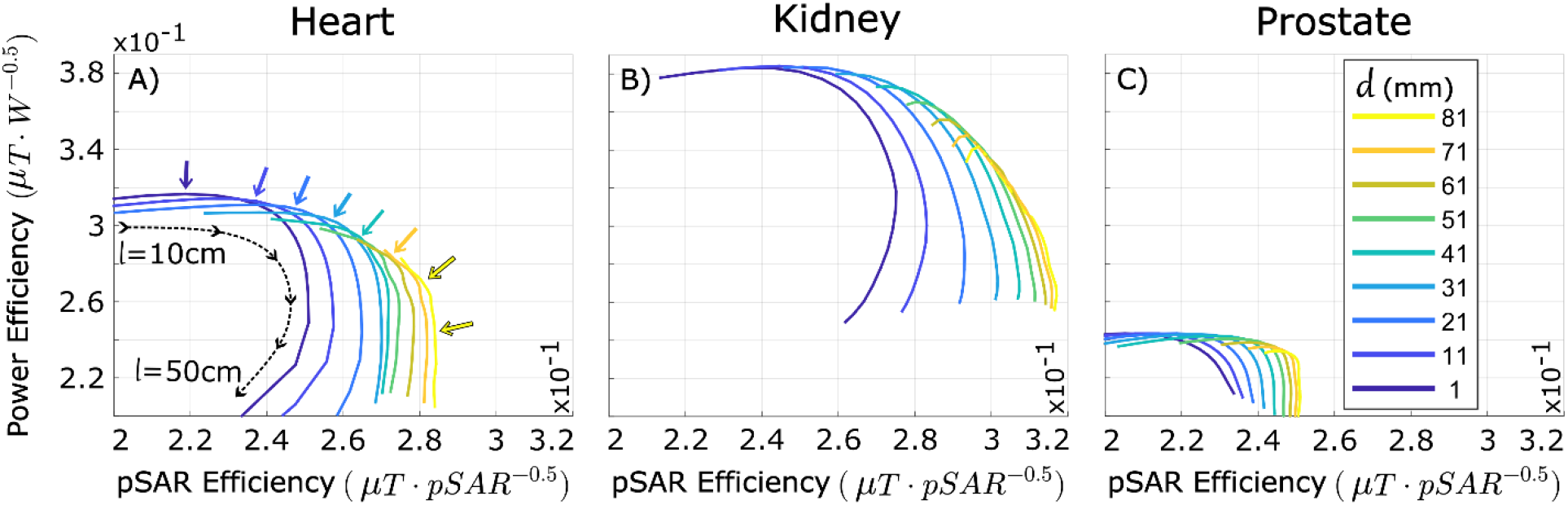
L-curves in the power efficiency by pSAR efficiency by efficiency space in the three different anatomies. Each line contains of all of the unique dipole lengths tested (100 to 500mm, illustrated by black dotted line in A). Both power efficiency and pSAR efficiency are higher in the more superficial organs like the heart and kidney. As is shown in Figure 2, increasing length ‘*l’* and distance-to-load ‘*d’* improves pSAR efficiency. Interestingly, the dominating edge of these curves, indicated by the colored arrows in A, is comprised of different element lengths each at varying distance-to-load. It is shown particularly well for the heart but the pattern does hold for the kidney and slightly holds in the prostate. This implies that the optimal length and distance-to-load varies based on anatomical target and system power constraints.

Short dipoles were more sensitive to variable loading compared to longer dipoles (Figure 4). Dipoles shorter than 140mm were unable to maintain a match better than -6dB after moving the elements a single centimeter closer or further from the 40mm to which they were originally matched. The 500mm half wavelength dipole was able to maintain a match better than -10dB after varying distance-to-load by 20mm in each direction.

**Figure 4.**
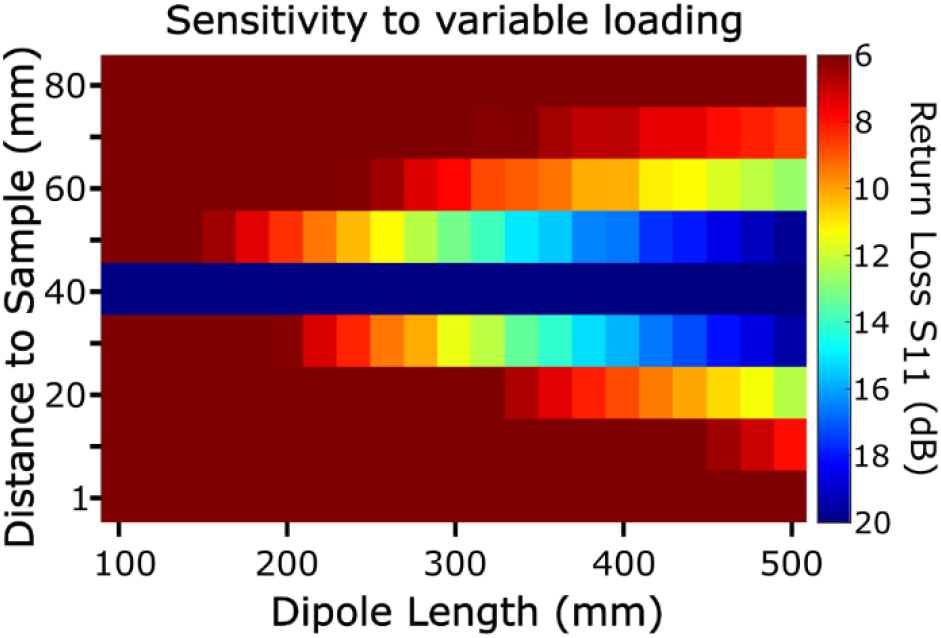
Element matching as a function of distance-to-load and length for the Duke kidney simulations. For each dipole length, the element was matched, via an L-match, to better than -20dB at *d* = 40mm. This L-match was then placed at the feed of all the other elements of the same length and the match at each distance-to-load was encoded as per the color bar. Short dipoles are extremely load dependent, with the match becoming worse than - 6dB when varying distance-to-load by only 10 mm. The half-wave dipole, at a length of 500mm, is the most robust to variations in distance-to-load.

The tradeoff between power-efficiency and pSAR-efficiency for meandered elements is shown in Figure 5A and followed a trend similar to the straight dipoles, with short dipoles close to the sample having the highest power efficiency and long dipoles further away having the highest pSAR efficiency. The larger meander width, *w*=80mm, had increased pSAR efficiency but decreased power efficiency. The addition of an RF-shield reduced pSAR efficiency while maintaining similar power efficiency (in Figure 5A, the solid lines of the same color are always to the right of the dashed lines of the same color). Figure 5B shows optimal homogeneity occurs at increased distance-to-load and element length while the addition of a shield and varying meander width had minimal impact. Performance of meandered and straight dipoles are shown together in Figures 5C and 5D which show meandered dipoles offer a more optimal balance of power efficiency, pSAR efficiency, and homogeneity.

**Figure 5.**
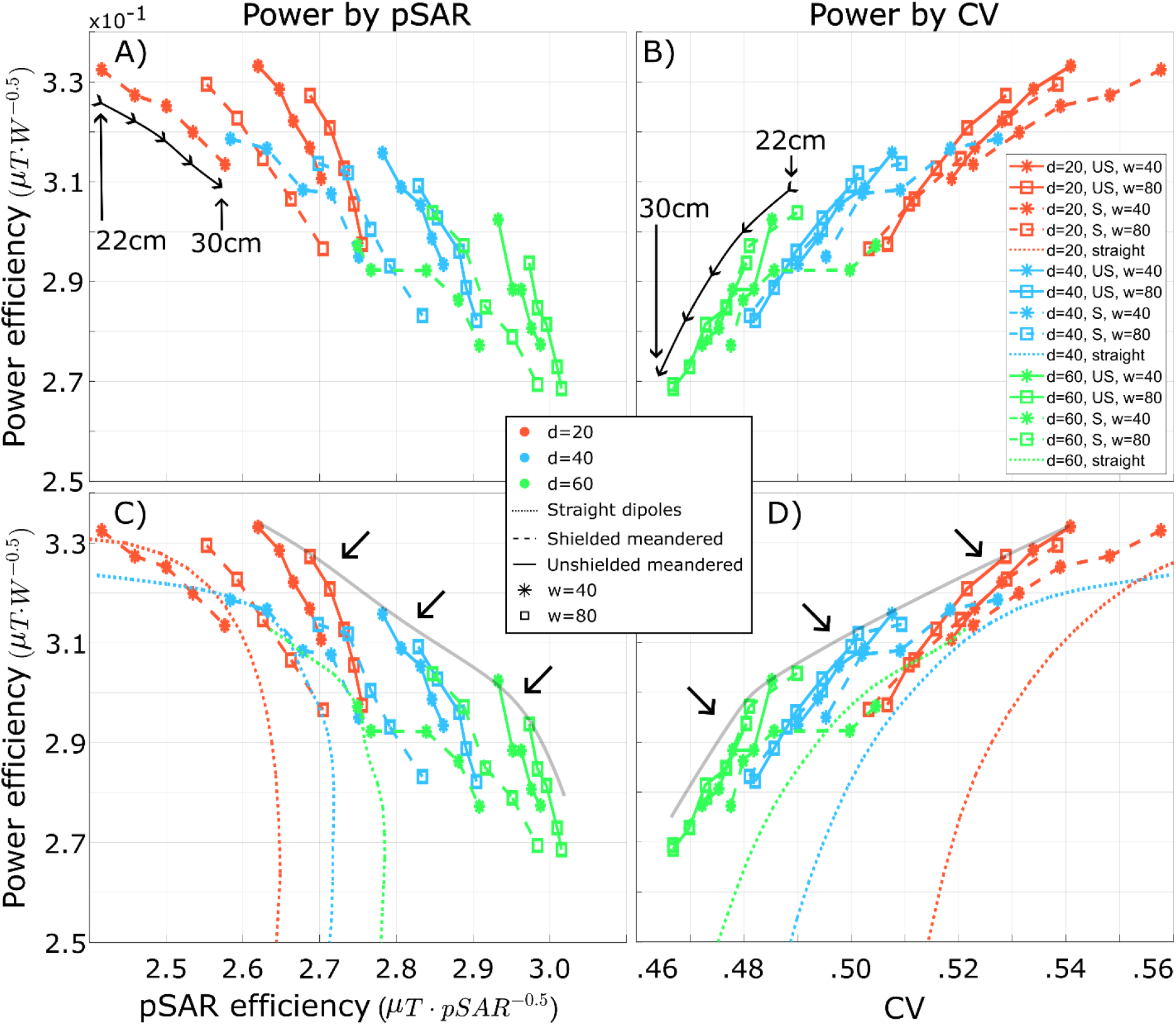
Meandered dipole performance plot in the power efficiency by pSAR efficiency space (A,C) and power efficiency by CV space (B,D). Color is used to encode distance-to-load ‘*d’*, markers encode width ‘*w’*, and line-style encodes element type. In subplots A and B, only the meandered dipole element performance is plotted. Each line in A and B contains the performance of 5 different meandered dipole lengths, varying from 220mm to 300mm in steps of 20mm (black line with arrows in A), with longer elements having better pSAR efficiency but lower power efficiency. Shielded and unshielded elements have nearly identical power efficiency but adding the shield decreased pSAR efficiency (compare dashed vs solid lines of the same color and symbol in A). The larger *w*=80mm meander had better pSAR efficiency but lower pSAR efficiency (compare square vs asterisk markers for each color and line type in A). Increasing *d* improved pSAR efficiency but decreased pSAR efficiency (compare the different colors for each marker and line type in A). The CV metric in B has tradeoffs similar to pSAR where longer dipoles at greater distances have a lower CV and therefore a more homogeneous field. Shielding and *w* had minimal impact on CV. Subplots C and D plot the straight dipole performance (unmarked dotted line) and meandered dipole performance. In these figures the unshielded meandered element performance now dominates straight dipole performance in both the power efficiency by pSAR efficiency and power efficiency by CV space (arrows in C and D pointing to the gray line indicates the dominating edge).

Experimental results were in good agreement with simulation results (Figure S1). Longer 500mm dipoles had the lowest power efficiency, 200mm dipole the highest power efficiency, and changing the distance-to-load from 10mm to 40mm reduced power efficiency but increased homogeneity as evidenced by the reduced overflipping on the surface.

### Array Coupling

Inter-element coupling was found to increase with increasing distance-to-load (Figure S2). Increasing element spacing, and shifting the elements in z reduced coupling in the array. The only simulation scenarios to achieve better than -10 dB decoupling at distances greater than 20mm occurred at *d* =40mm, *s*=120mm and z=60mm and 90mm.

## Discussion

The transition from remote transmitters at lower field strengths to local transmit arrays at 7T presents a unique set of challenges and design considerations. While the literature is rich with advancements in dipole geometry, a comprehensive analysis that investigates the coupled relationship between coil geometry and distance-to-load has not been presented. Our study addresses this gap by systematically evaluating power efficiency, pSAR efficiency, and homogeneity performance as a function of dipole geometry, lengths, distance-to-load, and with or without shielding.

The multi-objective nature of RF coil design is a critical consideration for clinical translation. Figure 3 reveals a fundamental trade-off between power efficiency and pSAR efficiency as distance-to-load and length are varied. This finding demonstrates that there is no single optimal distance-to-load, but rather a spectrum of choices that allow designers to balance performance metrics based on specific system constraints. For instance, a designer with ample RF power may choose an increased distance-to-load to maximize pSAR efficiency and homogeneity, while a system with limited power may opt for a shorter distance-to-load to boost power efficiency. Many 7T arrays up to this point have reduced distance-to-load in order to achieve a higher power efficiency, which could be attributed to the fact that the initial research-only 7T MRI systems were more power limited, often having around 8kW in total power available from the RFPAs. Conversely, when a remote 7T transmitter was constructed by Orzada et. al.^50^, the group utilized 32 1kW amplifiers in part to address the decreased power efficiency of the array. The multi-parameter, multi-objective analysis framework presented here provides a new, data-driven methodology for single-element optimization, making trade-offs between different performance metrics explicit and transparent.

A fundamental challenge in RF coil evaluation is the creation of meaningful performance metrics, which must contend with both a dimensionality mapping problem and the multi-objective nature of MRI. Because 3D EM fields must be distilled into scalar valued metrics, the definition of a metric such as “power efficiency” becomes ambiguous since it could be calculated as the maximum, minimum, or mean at some depth within a homogeneous phantom. The anatomical VOI based analysis presented in this paper addresses this ambiguity by calculating mean efficiencies and homogeneity over anatomically relevant volumes, providing a more intuitive and clinically relevant assessment. Furthermore, we presented a quantitative loading-sensitivity analysis showing that longer dipoles are more robust to variable loading.

By applying this multi-objective framework, this study provides a quantitative guide for RF coil designers to navigate the fundamental trade-offs between performance metrics. With known system specifications, designers can leverage the presented heatmaps and L-curves to identify optimal geometries and distances-to-load for a given application. A key finding is that the ∼20mm distance-to-load, common in many 7T body arrays^18–20,22,30,33^, yields good power efficiency but is suboptimal for pSAR efficiency and homogeneity. Our results suggest that increasing *d* to between 40 and 60 mm is a practical approach to improve pSAR efficiency. Furthermore, this larger distance would create valuable physical space for integrating local receive-only arrays, mirroring the dedicated transmit/receive configurations used at lower field strengths and prototype remote-transmit arrays at 7T^51^. Our analysis across different anatomies highlights the impact of evaluating performance in different anatomical targets. For instance, the heart, a large semi-superficial organ, exhibited significant homogeneity variations in the smaller distance-to-load and dipole length simulations which would translate into over-flipping near the transmit elements, an observation also reported at 3T when using local transitters^16^. This finding expands on the benefits to be gained from increasing distance-to-load by improving image quality in targets such as the heart. An important extension of this work will be to verify if these new VOI-based metrics are indeed better predictors of multi-element array performance compared to traditional metrics derived from 1D line projections.

While this study provides a relatively comprehensive analysis of dipole performance, it is subject to several limitations. We did not include an analysis of other common coil geometries, such as loop elements, as a function of distance-to-load. Furthermore, while the metrics reported for single-elements are expected to be positively correlated with array performance, the exploration of how these findings translate to an array configuration was outside the scope of this work. In our coupling analysis, dedicated decoupling networks^50,52^ were not included. Finally, there may exist naturally decoupled coil geometries^41,53,54^ for specific configurations that were not explored.

## Conclusion

To more fully utilize the fleet of 7T scanners currently installed around the world, body imaging needs to obtain regulatory approval. Providing a more user-friendly imaging setup with consistent and reproducible performance is pivotal to obtaining approval. A critical aspect of meeting both of these goals is the development of an optimized body imaging array. To build such an array, RF engineers need to fully understand how the many design degrees of freedom impact *all* the performance metrics of an RF coil. This study provides an in-depth analysis of many different dipole geometries, at variable distance-to-load, on different anatomical targets. We present new VOI based performance metrics and report these metrics simultaneously to illustrate the tradeoffs between power efficiency, pSAR efficiency, and homogeneity. These metrics are displayed individually in heatmaps and in the coupled power efficiency by pSAR efficiency space which demonstrates that there is a fundamental tradeoff between power efficiency and pSAR efficiency. Furthermore, a qualitative analysis on loading sensitivity was presented which demonstrated that longer dipole elements were more robust to variable loading conditions, a critical performance metric for reliable clinical imaging with variable patient body types. By quantifying the impact of both distance-to-load and length, this work provides a guide for designing optimal transmitter arrays while simultaneously establishing critical mechanical specifications for a potential integrated receive array, ensuring maximal performance of next-generation transmit/receive UHF body coils.

## REFERENCES

1. Pohmann R, Speck O, Scheffler K. Signal-to-Noise Ratio and MR Tissue Parameters in Human Brain Imaging at 3, 7, and 9.4 Tesla Using Current Receive Coil Arrays. doi:10.1002/mrm.25677

2. Wiesinger F, Van De Moortele PF, Adriany G, De Zanche N, Ugurbil K, Pruessmann KP. Parallel imaging performance as a function of field strength - An experimental investigation using electrodynamic scaling. Magn Reson Med. 2004;52(5):953–964. doi:10.1002/MRM.20281

3. Le Ster C, Grant A, Van de Moortele PF, et al. Magnetic field strength dependent SNR gain at the center of a spherical phantom and up to 11.7T. Magn Reson Med. 2022;88(5):2131–2138. doi:10.1002/MRM.29391

4. Guérin B, Villena JF, Polimeridis AG, et al. The ultimate signal-to-noise ratio in realistic body models. Magn Reson Med. 2017;78(5):1969–1980. doi:10.1002/mrm.26564

5. Ebrahimi B, Gloviczki M, Woollard JR, Crane JA, Textor SC, Lerman LO. Compartmental Analysis of Renal BOLD MRI Data: Introduction and Validation. Invest Radiol. 2012;47(3):175. doi:10.1097/RLI.0B013E318234E75B

6. Vink EE, De Boer A, Hoogduin HJM, et al. Renal BOLD-MRI relates to kidney function and activity of the renin-angiotensin-aldosterone system in hypertensive patients. J Hypertens. 2015;33(3):597–604. doi:10.1097/HJH.0000000000000436

7. Philips BWJ, Fortuin AS, Orzada S, Scheenen TWJ, Maas MC. High resolution MR imaging of pelvic lymph nodes at 7 Tesla. Magn Reson Med. 2017;78(3):1020–1028. doi:10.1002/MRM.26498

8. Frischer JM, Göd S, Gruber A, et al. Susceptibility-weighted imaging at 7 T: Improved diagnosis of cerebral cavernous malformations and associated developmental venous anomalies. Neuroimage Clin. 2012;1(1):116–120. doi:10.1016/J.NICL.2012.09.005

9. Li X, Auerbach EJ, Van de Moortele P, Ugurbil K, Metzger GJ. Quantitative single breath-hold renal arterial spin labeling imaging at 7T. Magn Reson Med. 2018;79(2):815–825. doi:10.1002/mrm.26742

10. Kraff O, Fischer A, Nagel AM, Mönninghoff C, Ladd ME. MRI at 7 Tesla and above: Demonstrated and potential capabilities. Journal of Magnetic Resonance Imaging. 2015;41(1):13–33. doi:10.1002/JMRI.24573

11. Georgakis IP, Polimeridis AG, Lattanzi R. A formalism to investigate the optimal transmit efficiency in radiofrequency shimming. NMR Biomed. 2020;33(11). doi:10.1002/NBM.4383

12. Fiedler TM, Ladd ME, Bitz AK. SAR Simulations & Safety. Neuroimage. 2018;168:33–58. doi:10.1016/j.neuroimage.2017.03.035

13. Orzada S, Akash S, Fiedler TM, Kratzer FJ, Ladd ME. An investigation into the dependence of virtual observation point-based specific absorption rate calculation complexity on number of channels. Magn Reson Med. 2023;89(1):469–476. doi:10.1002/MRM.29434

14. Eichfelder G, Gebhardt M. Local specific absorption rate control for parallel transmission by virtual observation points. Magn Reson Med. 2011;66(5):1468–1476. doi:10.1002/mrm.22927

15. Orzada S, Fiedler TM, Ladd ME. Hybrid algorithms for SAR matrix compression and the impact of post-processing on SAR calculation complexity. Magn Reson Med. 2024;92(6):2696–2706. doi:10.1002/MRM.30235

16. Weinberger O, Winter L, Dieringer MA, et al. Local Multi-Channel RF Surface Coil versus Body RF Coil Transmission for Cardiac Magnetic Resonance at 3 Tesla: Which Configuration Is Winning the Game? Published online 2016. doi:10.1371/journal.pone.0161863

17. Fiedler TM, Ladd ME, Orzada S. Local and whole-body SAR in UHF body imaging: Implications for SAR matrix compression. Magn Reson Med. 2025;93(2):842–849. doi:10.1002/MRM.30306

18. Ertürk MA, Raaijmakers AJE, Adriany G, Uğurbil K, Metzger GJ. A 16-channel combined loop-dipole transceiver array for 7 Tesla body MRI. Magn Reson Med. 2017;77(2):884–894. doi:10.1002/MRM.26153

19. Haluptzok TD, Lagore RL, Schmidt S, Metzger GJ. A shielded 32-channel body transceiver array with integrated electronics for 7 T. Magn Reson Med. 2025;94(2):852–866. doi:10.1002/MRM.30498

20. Schmidt S, Ertürk MA, He X, Haluptzok T, Eryaman Y, Metzger GJ. Improved 1H body imaging at 10.5 T: Validation and VOP-enabled imaging in vivo with a 16-channel transceiver dipole array. Magn Reson Med. 2024;91(2):513–529. doi:10.1002/MRM.29866

21. Zivkovic I, de Castro CA, Webb A. Design and characterization of an eight-element passively fed meander-dipole array with improved specific absorption rate efficiency for 7 T body imaging. NMR Biomed. 2019;32(8). doi:10.1002/NBM.4106

22. Raaijmakers AJE, Italiaander M, Voogt IJ, et al. The fractionated dipole antenna: A new antenna for body imaging at 7 Tesla. Magn Reson Med. 2016;75(3):1366–1374. doi:10.1002/mrm.25596

23. Orzada S, Bitz AK, Johst S, et al. Analysis of an Integrated 8-Channel Tx/Rx Body Array for Use as a Body Coil in 7-Tesla MRI. Front Phys. 2017;5(JUN):259150. doi:10.3389/fphy.2017.00017

24. Lattanzi R, Sodickson DK. Ideal current patterns yielding optimal signal-to-noise ratio and specific absorption rate in magnetic resonance imaging: Computational methods and physical insights. Magn Reson Med. 2012;68(1):286–304. doi:10.1002/mrm.23198

25. Raaijmakers AJE, Luijten PR, van den Berg CAT. Dipole antennas for ultrahigh-field body imaging: a comparison with loop coils. NMR Biomed. 2016;29(9):1122–1130. doi:10.1002/nbm.3356

26. Giannakopoulos II, Georgakis IP, Sodickson DK, Lattanzi R. Computational methods for the estimation of ideal current patterns in realistic human models. Magn Reson Med. 2024;91(2):760–772. doi:10.1002/MRM.29864

27. Pfrommer A, Henning A. The ultimate intrinsic signal-to-noise ratio of loop- and dipole-like current patterns in a realistic human head model. Magn Reson Med. 2018;80(5):2122–2138. doi:10.1002/MRM.27169

28. Orzada S, Bahr A, Bolz T. A novel 7 T microstrip element using meanders to enhance decoupling. In: Proc. Intl. Soc. Mag. Reson. Med. 16. 2008:2979.

29. Chen G, Cloos M, Sodickson D, Wiggins G. A 7T 8 channel transmit-receive dipole array for head imaging: dipole element and coil evaluation. In: Proc. Intl. Soc. Mag. Reson. Med. 22. 2014.

30. Steensma B, van de Moortele P, Ertürk A, et al. Introduction of the snake antenna array: Geometry optimization of a sinusoidal dipole antenna for 10.5T body imaging with lower peak SAR. Magn Reson Med. 2020;84(5):2885–2896. doi:10.1002/mrm.28297

31. Destruel A, Jin J, Weber E, et al. Integrated Multi-Modal Antenna with Coupled Radiating Structures (I-MARS) for 7T pTx Body MRI. IEEE Trans Med Imaging. 2022;41(1):39–51. doi:10.1109/TMI.2021.3103654,

32. Lagore RL, Moeller S, Zimmermann J, et al. An 8-dipole transceive and 24-loop receive array for non-human primate head imaging at 10.5 T. NMR Biomed. 2021;34(4). doi:10.1002/NBM.4472

33. Sadeghi-Tarakameh A, Adriany G, Metzger GJ, et al. Improving radiofrequency power and specific absorption rate management with bumped transmit elements in ultra-high field MRI. Magn Reson Med. 2020;84(6):3485–3493. doi:10.1002/MRM.28382

34. Sadeghi-Tarakameh A, Jungst S, Lanagan M, et al. A nine-channel transmit/receive array for spine imaging at 10.5 T: Introduction to a nonuniform dielectric substrate antenna. Magn Reson Med. 2022;87(4):2074–2088. doi:10.1002/MRM.29096,

35. Raaijmakers AJE, Ipek O, Klomp DWJ, et al. Design of a radiative surface coil array element at 7 T: The single-side adapted dipole antenna. Magn Reson Med. 2011;66(5):1488–1497. doi:10.1002/MRM.22886

36. van Leeuwen CC, Steensma BR, Klomp DWJ, van den Berg CAT, Raaijmakers AJE. The Coax Dipole: A fully flexible coaxial cable dipole antenna with flattened current distribution for body imaging at 7 Tesla. Magn Reson Med. 2022;87(1):528–540. doi:10.1002/MRM.28983

37. Budé LMI, Steensma BR, Zivkovic I, Raaijmakers AJE. The coax monopole antenna: A flexible end-fed antenna for ultrahigh field transmit/receive arrays. Magn Reson Med. 2024;92(1):361–373. doi:10.1002/MRM.30036

38. Deniz CM, Vaidya M V., Sodickson DK, Lattanzi R. Radiofrequency energy deposition and radiofrequency power requirements in parallel transmission with increasing distance from the coil to the sample. Magn Reson Med. 2016;75(1):423–432. doi:10.1002/mrm.25646

39. Hurshkainen AA, Steensma B, Glybovski SB, et al. A parametric study of radiative dipole body array coil for 7 Tesla MRI. Photonics Nanostruct. 2020;39:100764. doi:10.1016/J.PHOTONICS.2019.100764

40. Fiedler TM, Orzada S, Flöser M, et al. Performance analysis of integrated RF microstrip transmit antenna arrays with high channel count for body imaging at 7 T. NMR Biomed. 2021;34(7). doi:10.1002/NBM.4515

41. Yan X, Gore JC, Grissom WA. Self-decoupled radiofrequency coils for magnetic resonance imaging. Nat Commun. 2018;9(1):3481. doi:10.1038/s41467-018-05585-8

42. Rietsch SHG, Orzada S, Bitz AK, Gratz M, Ladd ME, Quick HH. Parallel transmit capability of various RF transmit elements and arrays at 7T MRI. Magn Reson Med. 2018;79(2):1116–1126. doi:10.1002/mrm.26704

43. Hubmann MJ, Orzada S, Kowal R, Anton Grimm J, Speck O, Maune H. Towards Large Diameter Transmit Coils for 7-T Head Imaging: A Detailed Comparison of a Set of Transmit Element Design Concepts. NMR Biomed. 2025;38(5):1–19. doi:10.1002/nbm.70030

44. Ruytenberg T, Webb A, Zivkovic I. Shielded-coaxial-cable coils as receive and transceive array elements for 7T human MRI. Magn Reson Med. 2020;83(3):1135–1146. doi:10.1002/MRM.27964

45. Avdievich NI, Solomakha G, Ruhm L, Henning A, Scheffler K. Unshielded bent folded-end dipole 9.4 T human head transceiver array decoupled using modified passive dipoles. Magn Reson Med. 2021;86(1):581–597. doi:10.1002/MRM.28711

46. Christ A, Kainz W, Hahn EG, et al. The Virtual Family—development of surface-based anatomical models of two adults and two children for dosimetric simulations. Phys Med Biol. 2010;55(2):N23–N38. doi:10.1088/0031-9155/55/2/N01

47. Gosselin MC, Neufeld E, Moser H, et al. Development of a new generation of high-resolution anatomical models for medical device evaluation: the Virtual Population 3.0. Phys Med Biol. 2014;59(18):5287. doi:10.1088/0031-9155/59/18/5287

48. Carluccio G, Erricolo D, Oh S, Collins CM. An approach to rapid calculation of temperature change in tissue using spatial filters to approximate effects of thermal conduction. IEEE Trans Biomed Eng. 2013;60(6):1735–1741. doi:10.1109/TBME.2013.2241764

49. Yarnykh VL. Actual flip-angle imaging in the pulsed steady state: A method for rapid three-dimensional mapping of the transmitted radiofrequency field. Magn Reson Med. 2007;57(1):192–200. doi:10.1002/MRM.21120

50. Orzadaid S, Solbach K, Gratz M, et al. A 32-channel parallel transmit system add-on for 7T MRI. Published online 2019. doi:10.1371/journal.pone.0222452

51. Rietsch SHG, Brunheim S, Orzada S, et al. Development and evaluation of a 16-channel receive-only RF coil to improve 7T ultra-high field body MRI with focus on the spine. Magn Reson Med. 2019;82(2):796–810. doi:10.1002/MRM.27731

52. Zivkovic I. Interelement Decoupling Strategies at UHF MRI. Front Phys. 2021;9:717369. doi:10.3389/fphy.2021.717369

53. Rietsch SHG, Quick HH, Orzada S. Impact of different meander sizes on the RF transmit performance and coupling of microstrip line elements at 7 T. Med Phys. 2015;42(8):4542–4552. doi:10.1118/1.4923177

54. Zhang B, Sodickson DK, Cloos MA. A high-impedance detector-array glove for magnetic resonance imaging of the hand. Nat Biomed Eng. 2018;2(8):570–577. doi:10.1038/S41551-018-0233-Y

55. Wu X, Tian J, Schmitter S, Vaughan JT, Ugurbil K, Van De Moortele PF. Distributing coil elements in three dimensions enhances parallel transmission multiband RF performance: A simulation study in the human brain at 7 Tesla. Magn Reson Med. 2016;75(6):2464–2472. doi:10.1002/MRM.26194

