## Supplementary material for "Distant Dipoles: A Multi-Parameter and Multi-Objective Analysis of RF Coil Performance For 7T Body MRI": Figure S1

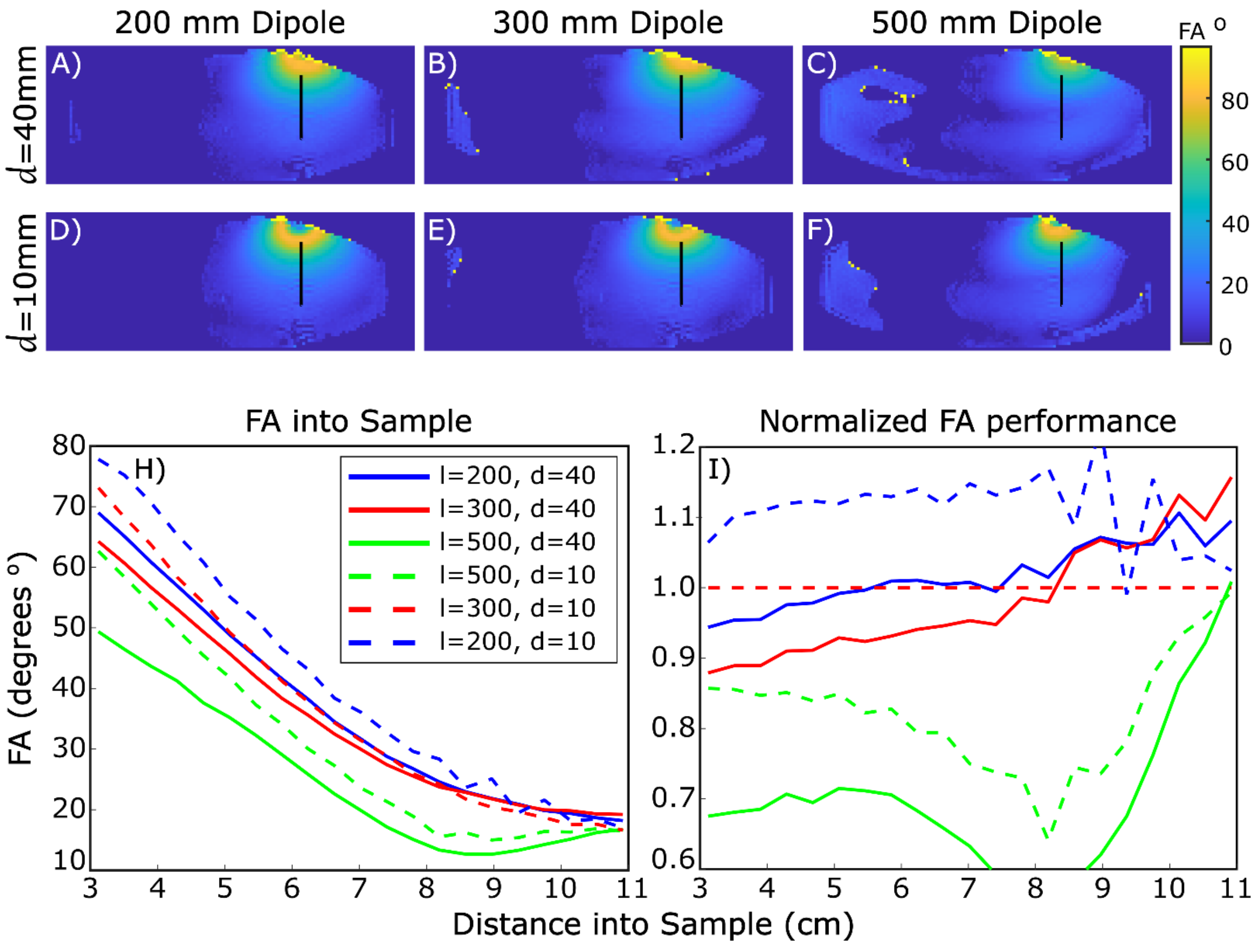


**Figure S1** Experimental AFI measurements (A-F) for 3 different dipole lengths at *d*=10mm and *d*=40mm. All six elements were able to be matched better than -18dB using a series inductor and a lumped element lattice balun. Flip angle profiles along the black lines in A-F are plot in H and normalized to the *l*=300mm, *d*=10mm configuration in subplot I. Experimental data has a trend similar to the simulation data with shorter and closer elements having better power efficiency. Interestingly, at greater depths into the phantom, the larger *d*=40mm configurations had greater power efficiency compared to the same element lengths at *d*=10mm. Furthermore, at a *d* of 40mm there was minimal overflipping on the surface which could point to increased homogeneity. Abbreviation: FA, flip angle; *d*, distance-to-load; *l*, dipole length.
