## Supplementary material for "Distant Dipoles: A Multi-Parameter and Multi-Objective Analysis of RF Coil Performance For 7T Body MRI": Figure S2

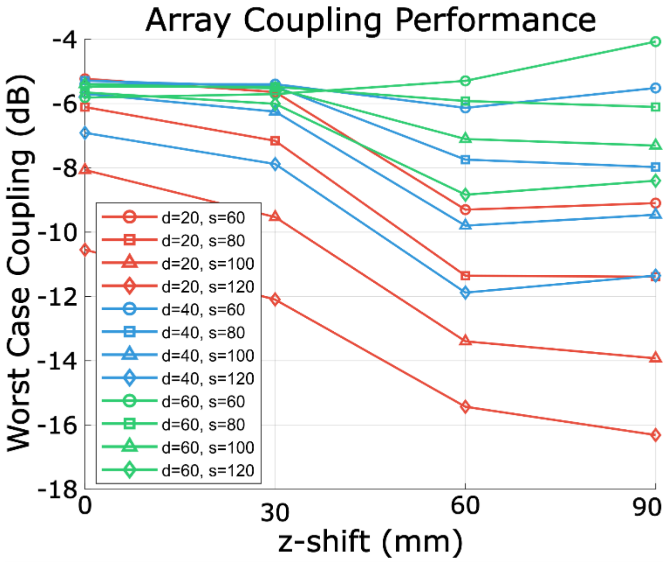


**Figure S2:** Coupling performance of a dipole array using 220mm long dipoles with variable distance-to-load '*d'*, element spacing '*s*', and z-shift. Shifting the elements in the z direction was found to help to decouple the elements. This decoupling, along with previously reported improvements in simultaneous multi-slice imaging performance,[55] suggests that z-shift is a parameter worthy of experimentation when designing transmitter arrays.
